# Molecular Basis of Sex Difference in Neuroprotection induced by Hypoxia Preconditioning in Zebrafish

**DOI:** 10.1101/2020.06.10.144097

**Authors:** Tapatee Das, Kalyani Soren, Mounica Yerasi, Avijeet Kamle, Arvind Kumar, Sumana Chakravarty

## Abstract

Hypoxia, the major cause of ischemia, leads to debilitating disease in infants via birth asphyxia and cerebral palsy, whereas in adults via heart attack and stroke. A widespread, natural protective phenomenon termed ‘Hypoxic Preconditioning’ occurs when prior exposures to hypoxia eventually results in robust hypoxia resistance. Accordingly, we have developed a novel model of sex-specific hypoxic preconditioning in adult zebrafish to mimic the tolerance of mini stroke(s) in human, which appears to protect against the severe damage inflicted by a major stroke event. Remarkable difference in the progression pattern of neuroprotection between preconditioning hypoxia followed by acute hypoxia (PH) group, and acute hypoxia (AH) group were observed with noticeable sex difference. Since gender difference has been reported in stroke, it was pertinent to investigate whether any such sex difference also exists in PH’s protective mechanism against acute ischemic stroke. In order to elucidate the neural molecular mechanisms behind sex difference in neuroprotection induced by PH, a high throughput proteomics approach utilizing iTRAQ was performed, followed by protein enrichment analysis using Ingenuity Pathway Analysis. Out of thousands of altered proteins in zebrafish brain, the ones having critical role either in neuroglial proliferation/differentiation or neurotrophic functions, were validated by analyzing their expression levels in PH, AH and normoxia groups. Results indicate that female zebrafish brains are more protected against the severity of AH. The study also sheds light on the involvement of many signaling pathways underlying sex difference in pre-conditioning induced neuroprotective mechanism, which can be further validated for the therapeutic approach.

## 1. INTRODUCTION

Hypoxic-ischemic injury is considered as one of the major causes of death and disability, worldwide. Interestingly, there is evidence that upon exposure to a hypoxic stimulus which is capable of causing injury up to a level less than the threshold to inflict damage endogenous protective mechanisms get activated and potentially attenuate the impact of the acute hypoxic attack that follows. Similarly, a sub-threshold hypoxic insult to an organ, has been shown to activate cellular pathways to reduce the damage caused by subsequent severe ischemic episodes - a phenomenon known as hypoxic preconditioning (HP) or hypoxic tolerance (HT) (Dirnagl *et al*., 2003). These observations suggest that HP is an adaptive reaction to a potentially noxious stimulus such as ischemia, hypoxia, hypoglycemia, or inflammation.

For maintaining the brain’s viability and proper functioning, uninterrupted oxygen supply is must. In fact, the entire central nervous system (CNS) is highly sensitive to changes in oxygen concentration due to high oxygen consumption (Luo *et al*., 2011). Therefore, during hypoxic episodes the brain cells or neural cells utilize key adaptive mechanisms that allow it to survive and maintain homeostasis. The plasticity nature of the brain is the major focus of HP research as it confers on lifelong ability to adapt based on the external or internal environmental challenges. With subsequent hypoxic insults the neural cells develop hypoxic tolerance which stimulates neuroplasticity with enhanced homeostatic control and the phenomenon is called modulation. During HP, modulation simultaneously works with plasticity and functions to sustain it. Based on the significance clinically novel therapeutics can be designed based on the mechanisms underlying plasticity and modulation (Lu *et al*., 1999b; Lu *et al*., 2005; Wang *et al*., 2007). The phenomenon of adaptation to hypoxia has been rigorously studied so far. For example, many studies have been done on acclimation to high altitude and the effects of chronic hypoxia. This paved the way for a base to understand the physiological responses to hypoxia at system level. However, the adaptive response to hypoxia may not be identical as the conditions like time and severity of hypoxia in these settings differ from that in pathological processes, therefore studying the cellular and molecular mechanisms of hypoxic tolerance may yield few critical targets for developing stroke therapeutics.

Adaptation to hypoxia at cellular level through the traditional knowledge based on systemic respiratory and cardiovascular responses, prior to HP term was introduced, Haldane named it as physicochemical brain (Haldane, 1927). An acquired tolerance of tissue-cells to hypoxia was thought to have developed evolutionary. Later, it was termed as “tissue-cell adaptation to hypoxia” (Michiels, 2004). Subsequently, animal models of hypoxia were developed to explore HP from the vantage point of behaviour, neurophysiology, neurochemistry, neuromorphology, and molecular biology.

This remarkable effect of HP as a mode of neuroprotection has been shown through many studies lately. In one of the study utilizing mice, there was significant increase of survival time in a hypobaric chamber when treated with brain homogenate from hypoxically preconditioned mice in comparison to a saline-treated control group (Lu *et al*., 1999a). Similarly, cells co-cultured under hypoxia with brain extract from hypoxically preconditioned animals were found to be substantially more viable than cells from the control group where no extract was added (Perez-Pinzon *et al*., 1996). The release of lactate dehydrogenase (LDH), an indicator of cell death, in cortical synaptosomes co-cultured with hypoxic preconditioned brain extract was progressively reduced, indicating protection by the extract. Another study indicated that histone acetylation is vital in conferring tolerance to brain hypoxia in rats exposed to hypobaric hypoxia (Samoilov *et al*., 2016). In a study that examined the effects of HP on white matter damage following a hypoxic-ischemic insult in neonatal rats, HP protected myelin, either directly or by enhancing oligodendrocyte progenitor maturation to replace damaged myelin (Suryana and Jones, 2014).These neurochemical effects of preconditioned brain extract suggest there are adaptive changes at the molecular level that confers tolerance to hypoxic stress. HP can be further simplified in relation to the time taken for adaptive response to the hypoxic stimulus. In fact, the efficacy of preconditioning protection to robust ischemic condition seems to be dependent on the difference in the degree of hypoxia and the delay between preconditioning and ischemia (Miller *et al*., 2001). Rapid HP, a protective phenotype lasting for a shorter duration, is induced in an early phase of adaptation which lasts from minutes to several hours after sublethal hypoxic exposure (Perez-Pinzon *et al*., 1996; Schurr *et al*., 1986). This early adaptation is believed to be a consequence of alterations in ion channel permeability, protein phosphorylation, and other post-translational modifications (Li *et al*., 2017).

Contrary delayed HP is best studied as classical preconditioning gene activation and de novo protein synthesis. This is first demonstrated only hours to days following PC. In delayed PC, a diverse family of pro-survival genes and proteins that improve the brain-tolerance to ischemic insults, is activated. This involves both; inhibition of injury mechanisms and a subsequent stimulation of underlying survival and repair mechanisms (Stagliano *et al*., 1999).

The protective effect of HP has been explained in various animal species and in organs such as the brain, kidneys, heart and skeletal muscle (Azad *et al*., 2012). Despite several previous studies on hypoxia signalling in model organisms and in mammals, the molecular pathways responsible for the protective effect of HP remain poorly understood (Azad and Haddad, 2013; Chang and Bargmann, 2008; Wacker *et al*., 2012). In this study, we developed a novel *in vivo* model of hypoxic preconditioning in adult zebrafish to discover novel markers (genes/proteins) that contribute to HP-induced neuroprotection. In light of the reports where gender difference has been clearly observed in stroke risk factors and disease history, we tried to investigate whether any such sex difference also exists in HP’s protective mechanism against acute ischemic stroke. In the present study, unlike other validated preconditioning protocols from *in vitro* and *in vivo* rodent models where hypoxia had been given at subthreshold level just once prior to the final hypoxic insult, we have developed a protocol that mimics multiple mini stroke-like conditions in human. Here, male and female zebrafish were exposed to hypoxia intermittently for three months, before one final acute hypoxic attack. This was followed by behavioural characterization of animals and sex-specific molecular responses in brain using high throughput proteomic study. The outcome of the study sheds light on the effect of PH on the neuroprotective processes that involved changes in adult brain neural stem or progenitor cells (NSCs/NPCs) and their differentiation, i.e. adult neurogenesis, in addition to alteration in many signalling pathways.

## 2 MATERIALS AND METHODS

### 2.1 Animal husbandry experimental setup for hypoxia preconditioning

A total of 120 adult zebrafish (5-8 months old, 60 for each sex) from the TL/AB strain were used in this study. The animals were well maintained using the standard procedures (Westerfield, 2000) and divided into six groups (n=20 in each group) named Normoxia Male (NM), Normoxia female (NF), Hypoxia preconditioning male (PHM), Hypoxia preconditioning female (PHF), Acute hypoxia male (AHM) and Acute hypoxia female (AHF). Animals in all the groups were maintained in the same environment where hypoxia preconditioning group animals were given intermittent hypoxia treatments (DO=0.6-2.0 mg/l/ 5-10 minutes) for a period of three months. The final day before sacrificing the animals, the Acute hypoxia male/female group along with hypoxia preconditioning male/female group were subjected to severe acute hypoxia treatment (DO=0.6 mg/l for 5 minutes) following recovery in normal water (DO=7 mg/l for an hour). After recovery period all the groups of animals were taken for the behaviour test (NTT) and post 4 hour of acute hypoxia all the animals were sacrificed for all the molecular analysis as reported earlier (Das *et al.*, 2019).

### 2.2 Behaviour analysis

Behavioural evaluations were done by the Novel Tank Test. First the individual response was recorded with video camera (Sony HDR/CX200) and later the data were analysed offline by Ethovision XT 10. In each test several behavioural parameters were observed and obtained the quantified values. Average of each group was taken in every parameter and standard deviation and standard error of mean (SEM) were calculated accordingly.

### 2.3 Immunohistochemistry for assessing cell death and neural proliferation markers

The fixed brain sections from individuals of different groups (n=6) were processed for immunohistochemistry (IHC) with several antibodies. The protocol was followed as described in previous report (Das *et al*., 2019). The PFA embedded brain from all the six groups were sectioned (30 μm thickness) using a cryomicrotome (Leica). TUNEL assays were carried out using the ApopTag Detection Kit (Chemicon), according to the manufacturer’s instructions. In every experimental series, negative (omitting the TdT enzyme) control slides were included. Among other primary antibodies, neural proliferation and differentiation markers GFAP (AB 7260, 1:1000), BLBP (AB 32423, 1:1000), S100 beta (AB 52642, 1:1000), NEUN (MAB-377, 1: 1000) and Tuj1 (AB18207, 1: 1000) were used.

### 2.4 Protein extraction for iTRAQ

The animals were euthanatized and quickly decapitated to remove the brain. The whole brains were homogenized in lysis buffer [50 mM ammonium bicarbonate (pH 8.0), 0.1% SDS with protease inhibitor cocktail from (SIGMA)] and for further efficient disruption and homogenization of tissue, a mild sonication was done using Bioruptor. The obtained lysates were cold-centrifuged at 14000 g for 15 minutes, and the protein concentration was quantified in collected supernatant using Barford assay with BSA as a standard. Further protein sample was cleaned up by acetone precipitation. For each group 80 μg of protein was taken and six volume of chilled acetone was added for precipitation and samples were air dried after decanting acetone.

Prior to trypsin digestion all the protein samples were gone through reduction and cysteine blocking using the reagents provided with iTRAQ 4-plex kit (AB SCIEX). Digestion and labelling of proteins were done according to manufacturer’s protocol. Subsequently all the labelled samples were pooled, vacuum dried and further cleaned up using C18 desalting column (Pierce Thermoscientific). The final fraction again concentrated using vacuum concentrator and reconstituted with 10 μl of 0.1% formic acid for LC-MS/MS analysis.

### 2.5 LC-MS/MS Analysis

Tandem mass spectrometric analysis of peptide samples from six groups (NM, NF, AM, AF, PM and PF) were carried out using LTQ-Orbitrap Velos coupled with Proxeon easy nLC system (Thermo scientific, Germany). Peptide samples were enriched using a C18 trap column and separated on an analytical column at flow rate of 350 nl/min using a linear gradient of acetonitrile (ACN) over 90 min. Mass spectrometric analysis was carried out in a data dependent manner with full scans acquired using the Orbitrap mass analyser at a mass resolution of 60,000 at m/z 400. The most intense precursor ions were selected from each MS scan for fragmentation using higher energy collision dissociation (HCD) with 50% normalized collision energy. The lock mass option was enabled for accurate mass measurements and the entire procedure was repeated to generate triplicate run.

### 2.6 Bioinformatic and Protein enrichment analysis

The obtained raw files of each fraction and its triplicates were analysed using Proteome Discoverer (Thermo Fisher Scientific version1.4,). MS/MS search was carried out using SEQUEST search engine against the NCBI RefSeq zebrafish protein database. Search parameters included trypsin as an enzyme with 1 missed cleavage allowed; precursor and fragment mass tolerance were set to 10 ppm and 0.5 Da, respectively; Methionine oxidation was set as a dynamic modification while methylthio modification at cysteine and iTRAQ modification at N-terminus of the peptide were set as static modifications. The FDR was calculated by enabling the peptide sequence analysis using a decoy database. High confidence peptide identifications were obtained by setting a target FDR threshold of 1 % at the peptide level. Proteins identified from the triplicate runs were selected for its differential expression in both the test samples as AH and PH against normoxia samples. Proteins having more than 1-log and less than 0.5-log fold changes in the test samples against the control samples were listed as differentially regulated in the tissue.

To perform functional enrichment tests of the candidate proteins, Ingenuity Pathway Analysis (IPA) software was used for both canonical pathways and molecular networks. The IPA system provides a more comprehensive pathway resource based on manual collection and curation. The rich information returned by IPA is also suitable for pathway crosstalk analysis as it has more molecules and their connections included. For analysis we have provided the identified peptides with relative and absolute expression fold change values.

### 2.8 RNA isolation and qPCR

Total RNA isolation was carried out by Trizol reagent with manufacturer’s instruction. cDNA was synthesized using RevertAid H Minus First Strand cDNA Synthesis kit as reported [8]. Primer sequences and PCR conditions are available upon request. Quantitative real-time PCR (qPCR) was performed in triplicates by using SYBR Green PCR Master Mix Detection System (Applied Biosystems, USA). Relative gene expression analysis was performed with β-actin as housekeeping gene.

### 2.9 STATISTICAL ANALYSIS

Statistical analysis was performed using Prism 8 (GraphPad) and Microsoft Excel. Graphs for each parameter were plotted for the averages obtained and the error bars represented the SEM of the values. Mean differences between groups were appropriately determined by either a twotailed unpaired Student’s t-test with confidence interval of 95% or two-way analyses of variance (ANOVA) followed by Post-hoc comparisons were performed by Tukey’s test. Statistical significance was set at *p* value <0.05. All the results were expressed as mean ± S.E.M.

## 3 RESULTS

### 3.1 Analysis for sex difference in behaviour of zebrafish after Preconditioned hypoxia and Acute hypoxia

Note: NM/NF = Normoxia Male/Normoxia Female, maintained in normoxia environment (DO = ~7-8 mg/l); AHM/AHF= Acute Hypoxia Male/Acute Hypoxia Female, maintained in normoxia environment until acute hypoxic exposure; PHM/PHF = Preconditioned Hypoxia Male/Preconditioned Hypoxia Female, maintained most of the time in normoxia environment but intermittently administered hypoxic stress over 3 months-period, followed by acute hypoxic exposure along with the animals in AH group (PH = PH + AH).

The experimental procedure was described in Fig.1A. After one hour of acute hypoxia treatment, animals from all the four treatment groups (AHM, AHF, PHM and PHF) along with two control groups (NM and NF) were introduced to the Novel Tank Test (NTT) and their individual behavioural responses were video recorded. The behaviour data (n=20 in each group) were later analysed in offline mode using EthoVision XT 10, for the locomotor activity including vertical exploration in the tank. The hypoxic response was not uniform in two groups (only acute and preconditioned). Although two-way ANOVA analysis for total distance travelled [F (2, 86) = 2.096; p< 0.1292], velocity [F (2, 86) = 2.242; p< 0.1124] did show significant changes only in females, in lower to upper zone frequency [F (2, 86) = 6.431; p<0.0025] and meandering [F (2, 86) = 25.55; p<0.0001] significant changes were observed in both the sexes. In general, considerable improvement in locomotion and exploration were observed in both the sexes. Interestingly, on assessment of vertical exploration behaviour and meandering, a hallmark of hypoxia characteristic behaviour, preconditioning hypoxia groups were found to be very much protected after the severe hypoxia treatment as compare to acute hypoxia group. Statistical analysis of data showed significant difference in acute hypoxia and preconditioning hypoxia group in showing adaptation to stress phenomenon in preconditioning hypoxia group (Fig 1).

**Fig. 1:**
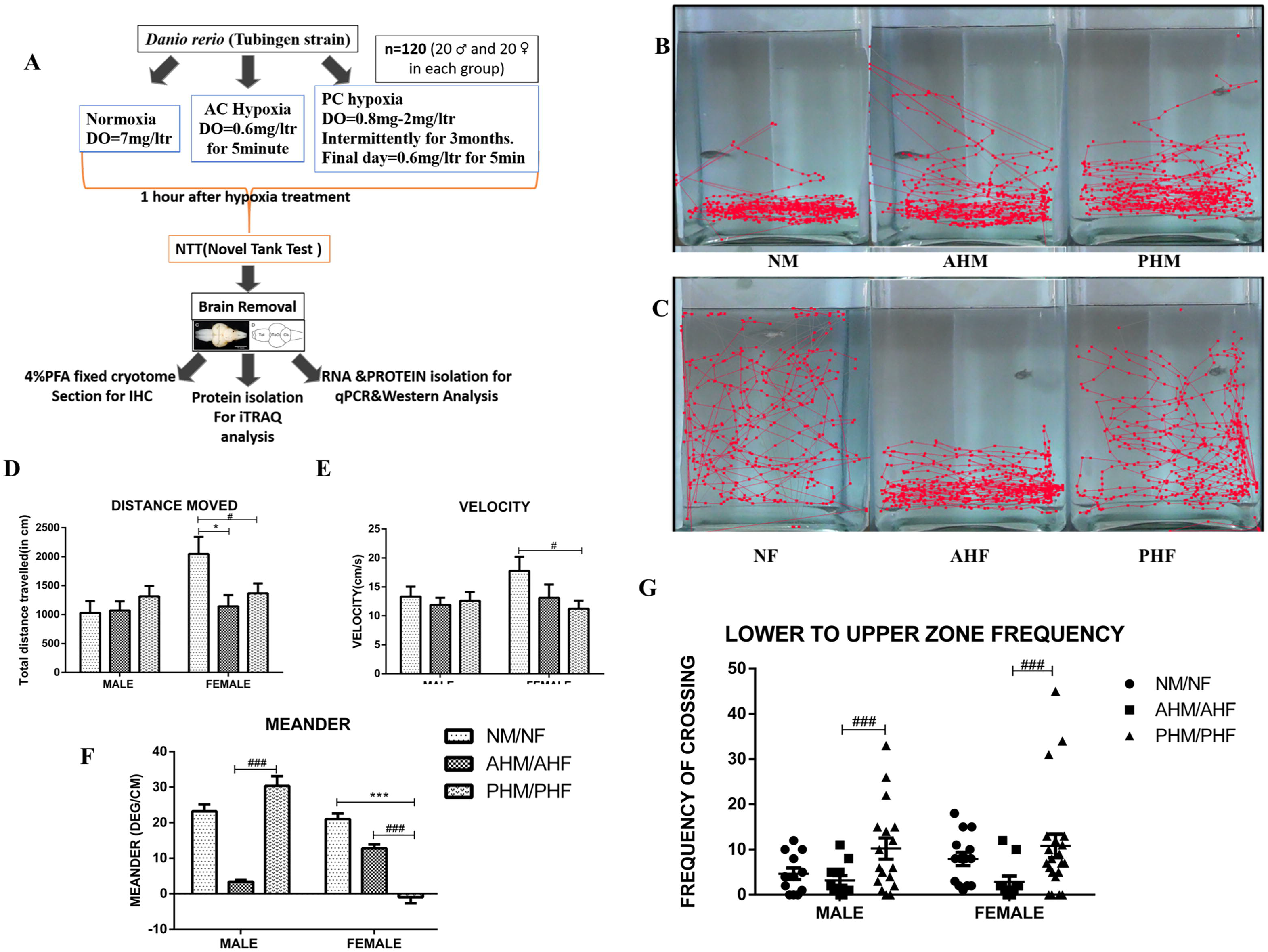
Sex-specific comparative analysis of neurobehaviour upon Acute hypoxia vs preconditioning hypoxia. Experimental workflow for studying neuro protection in preconditioning hypoxia (A). Represented track file of EthoVision XT for NM, AHM and PHM (B), NF, AHF and PHF (C). Analysis of basic locomotor parameters; total distance travelled (D), velocity (E), and meandering (F). Exploration of fish represented as lower to upper zone frequency in the novel tank (G). Data represent means □±□ SEM (\**p*□<□0.05), *n*□=□20 per group and data analysed by two-way ANOVA followed by Tukey post hoc analysis. * PHM /PHF vs NM /NF and # PHM/PHF vs AHM/AHF.

### 3.2 Evaluating sex difference in DNA damage in preconditioned hypoxia vs Acute hypoxia/ Normoxia

To evaluate the degree of neuroprotection in preconditioning hypoxia group we have performed TUNEL assay and analysed the data in comparison with acute hypoxia and normoxia group (n=15). A uniform protection against DNA damage in Preconditioning groups was observed in both sexes (Fig 2). As compared to normoxia group, acute hypoxia group showed a greater number of TUNEL positive cells where in preconditioning group the number of TUNEL positive cells are significantly less than acute hypoxia group. It is noteworthy to notice an indication of sex difference in DNA damage protection in preconditioning hypoxia group where PHF showed nearly equal number of TUNEL positive cells as NF unlike PHM/ NM combination. Comparison of the numbers of TUNEL-positive cells showed significant difference in two-way ANOVA analysis with [F (2, 62) =18.47; p<0.0001].

**Fig. 2:**
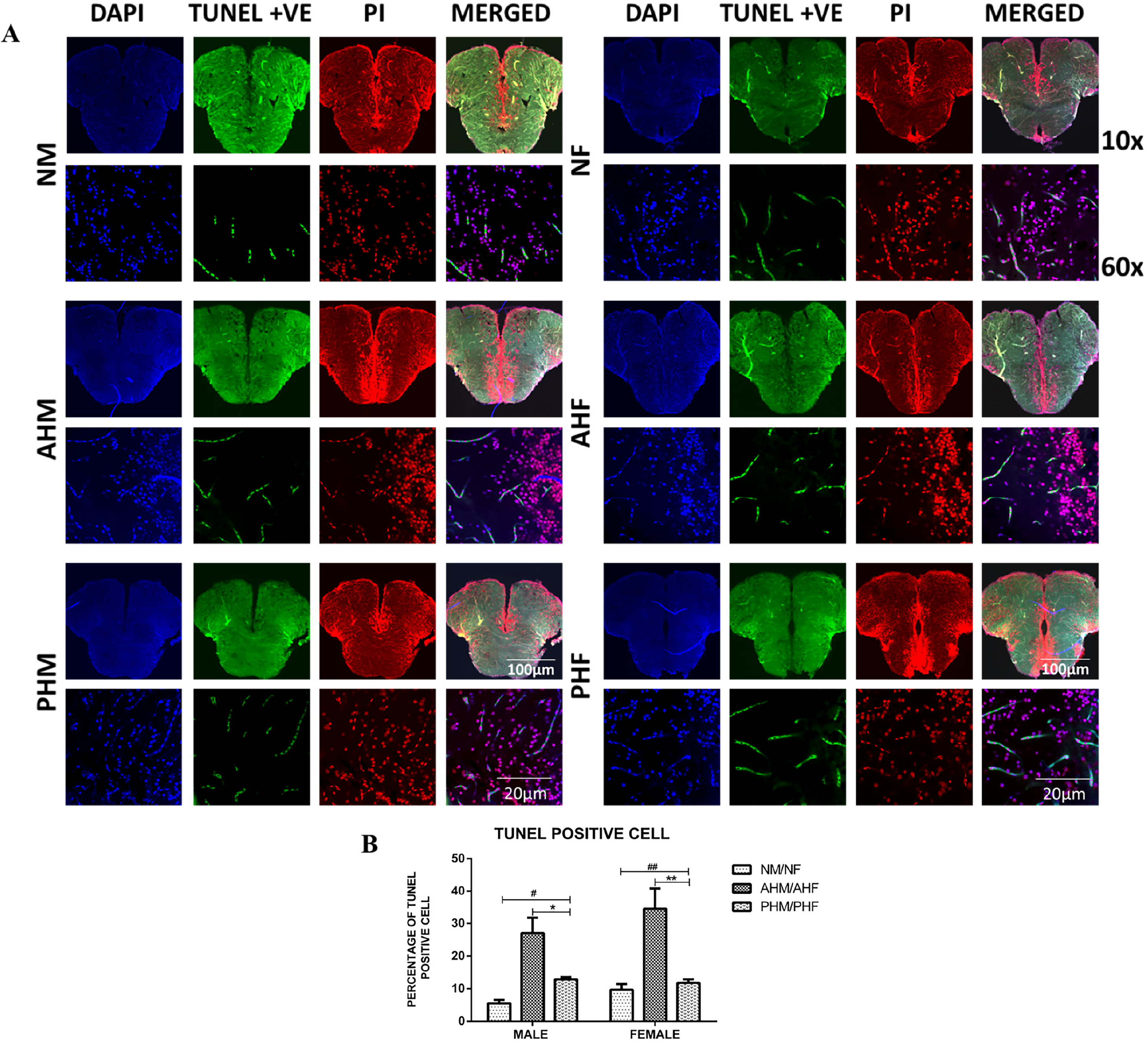
Sex-specific analysis of DNA damage protection in preconditioning hypoxia. TUNEL representative confocal images of 60x zebrafish brain sections (A). Bar graph indicates percentage of TUNEL positive cells (B), n=15. p <0.0001, * PHM /PHF vs NM /NF and # PHM/PHF vs AHM/AHF.

### 3.3 Quantitative brain proteome analysis by iTRAQ and protein enrichment analysis between Preconditioned hypoxia vs Acute hypoxia/ Normoxia

In this study, we have performed the triplicate runs of nLC-MS/MS analysis of iTRAQ labelled tryptic digests of proteins from male and female brain separately for Normoxia, Acute hypoxia, Preconditioned hypoxia groups. Differentially expressed proteins were identified from these fractions using iTRAQ and LC-MS/MS analysis on LTQ Orbitrap Velos mass spectrometer. The overall work flow of the study is shown in Fig 3A. Total 3017 proteins in female groups and 2581 proteins in male group were identified and quantified. In both male and female groups the expression of proteins were compared between PH/AH vs Normoxia for observing the effect of acute stress on both; with (PH) or without preconditioning (AH). A total of 2157 proteins were identified to be common in both male and female with varied expression status (Fig 3B). These common 2157 proteins based on their pattern of expression were shown through a heatmap (Fig 3C). Further the data were analysed based on their expression fold change. The values ranging from 0.1-0.7 were considered as downregulation and values more than 1.3 were considered as upregulation. In male among 2581 identified proteins, 2052 proteins in AHM vs NM and 2081 proteins PHM vs NM were upregulated whereas in female among 3017 proteins only 167 proteins in AHF vs NF and 72 proteins in PHF vs NF were upregulated. In male 213 proteins in AHM vs NM and 212 proteins in PHM vs NM were downregulated whereas in female 43 proteins in AHF vs NF and 36 proteins in PHF vs NF were downregulated. The data clearly indicated a noticeable sex difference in the altered expression levels of proteins. A major portion ~80% of the proteins in male were found upregulated and ~8 % of proteins were down regulated whereas in female only ~2-5 % of proteins were found upregulated and ~1.4 % of proteins were down regulated. Interestingly, where only ~12 % of proteins in male did not show any change in expression, in female most of the proteins [~92 %] remain unchanged by hypoxia. (Fig 3 C)

**Fig. 3:**
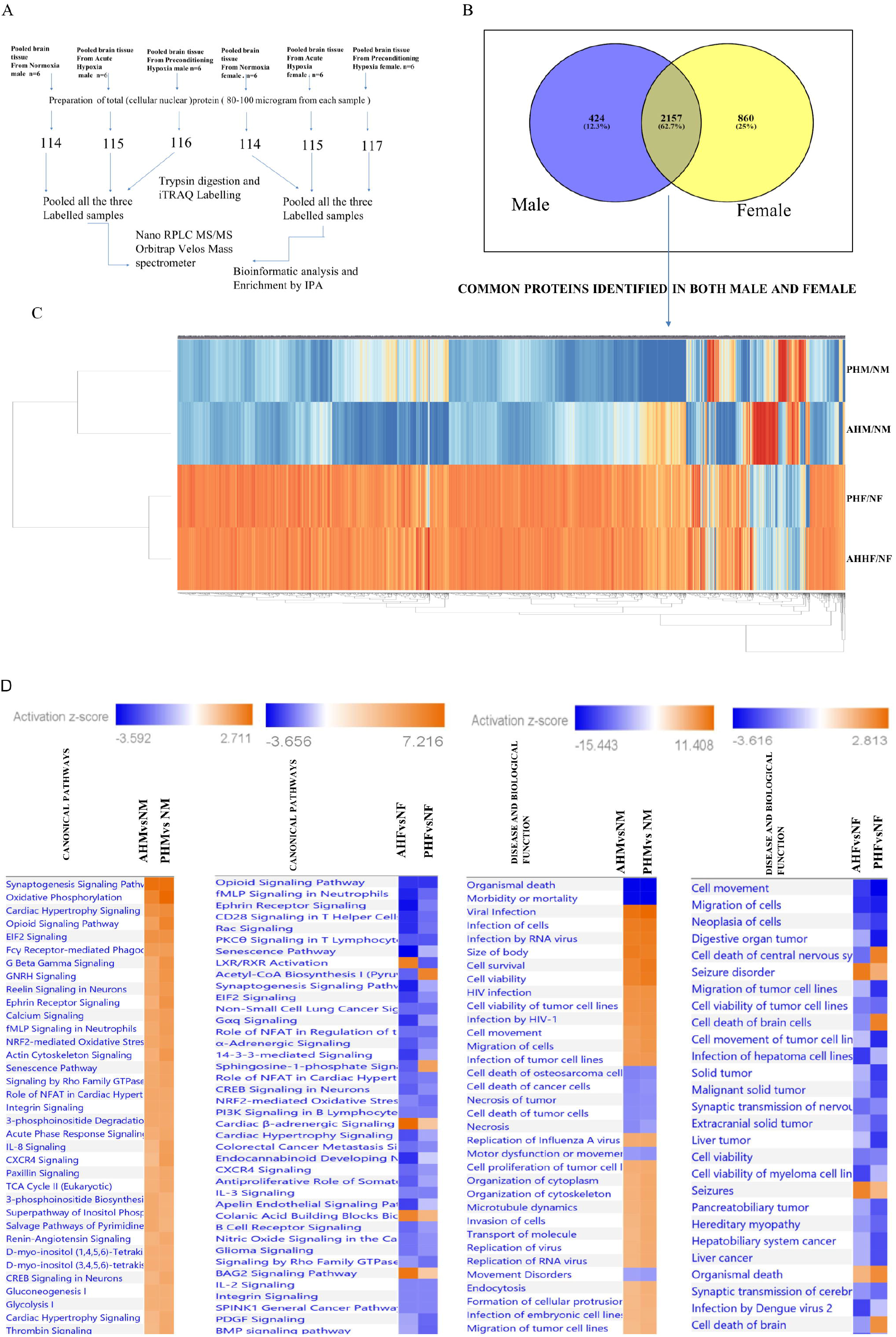
Sex-specific zebrafish brain proteome analysis in preconditioning hypoxia followed vs acute hypoxia by ITRAQ. Experimental workflow for iTRAQ (A). Venn diagram showing common proteins found in male and female brain proteome (B). Heat map for common proteins (2156) expression in AHM/F vs NM/F and PHM/F vs NM/F (C). IPA generated comparison heat map for canonical pathways and disease biological function (D).

The protein enrichment analysis by IPA based on its core expression analysis shed light on canonical pathways, upstream regulators, and disease function molecules altered in acute and preconditioning hypoxia groups. In both male and female, AH vs Normoxia and PH vs Normoxia were compared for the IPA core analysis. Out of the top five canonical pathways affected in male and female, four were found to be common in both (EIF2 Signalling, Mitochondrial Dysfunction, Synaptogenesis Signalling, Sirtuin signalling). But Oxidative Phosphorylation pathway was observed only in male animals, while the Regulation of eIF4 and p70S6K Signalling was observed only in females (Fig 3 D).Most of the canonical pathways obtained through core analysis for male were upregulated where in female downregulated(Fig 3 D). The comparison of canonical pathways in male for AHM vs NM and PHM vs NM showed not much contrasting type of regulation whereas in female AHF vs NF and PHF vs NF showed contrasting regulation in few canonical pathways (Fig 3 D). The upstream regulators identified through IPA for both male and female groups showed two distinct regulators, MYC and HTT, and many top analysis ready molecules for both the observations (observation 1 where AH vs NM/NF and observation 2 where PH vs NM/NF) were obtained through IPA mostly focuses on molecules with sex difference and AH vs PH as well.(Summary of IPA analysis report provided in supplementary). Although the protein enrichment analysis shed light on many novel targets from the top signalling pathways, due to the limitation of the commercially available zebrafish specific antibodies the validation of those targets was not done in the present study. Instead, we validated some other altered proteins in our data that are involved in neuroglial proliferation, differentiation, activation of growth factors, events underlying precondition hypoxia-induced neuroprotection.

### 3.4 Evaluating sex difference in neural cell death using apoptotic marker cleaved Caspase-3

Since cleaved Caspase −3 is known to be a strong marker of apoptotic cell death induction, here we mapped the expression of cleaved caspase-3 in brain by immunofluorescence chemistry (IFC) to assess the degree of protection against cell death in the preconditioning hypoxia group, compared to acute hypoxia and normoxia group (Fig 4). The percentage of caspase-3 positive cells in acute hypoxia group was found significantly more than that in normoxia group, whereas the preconditioning hypoxia group displayed protection by showing comparatively less percentage of caspase-3 positive cells than the acute hypoxia group. The two-way ANOVA showed significant result [F (2, 72) = 10.86; p<0.0001], as shown in graphical representation (Fig 4B).

**Fig. 4:**
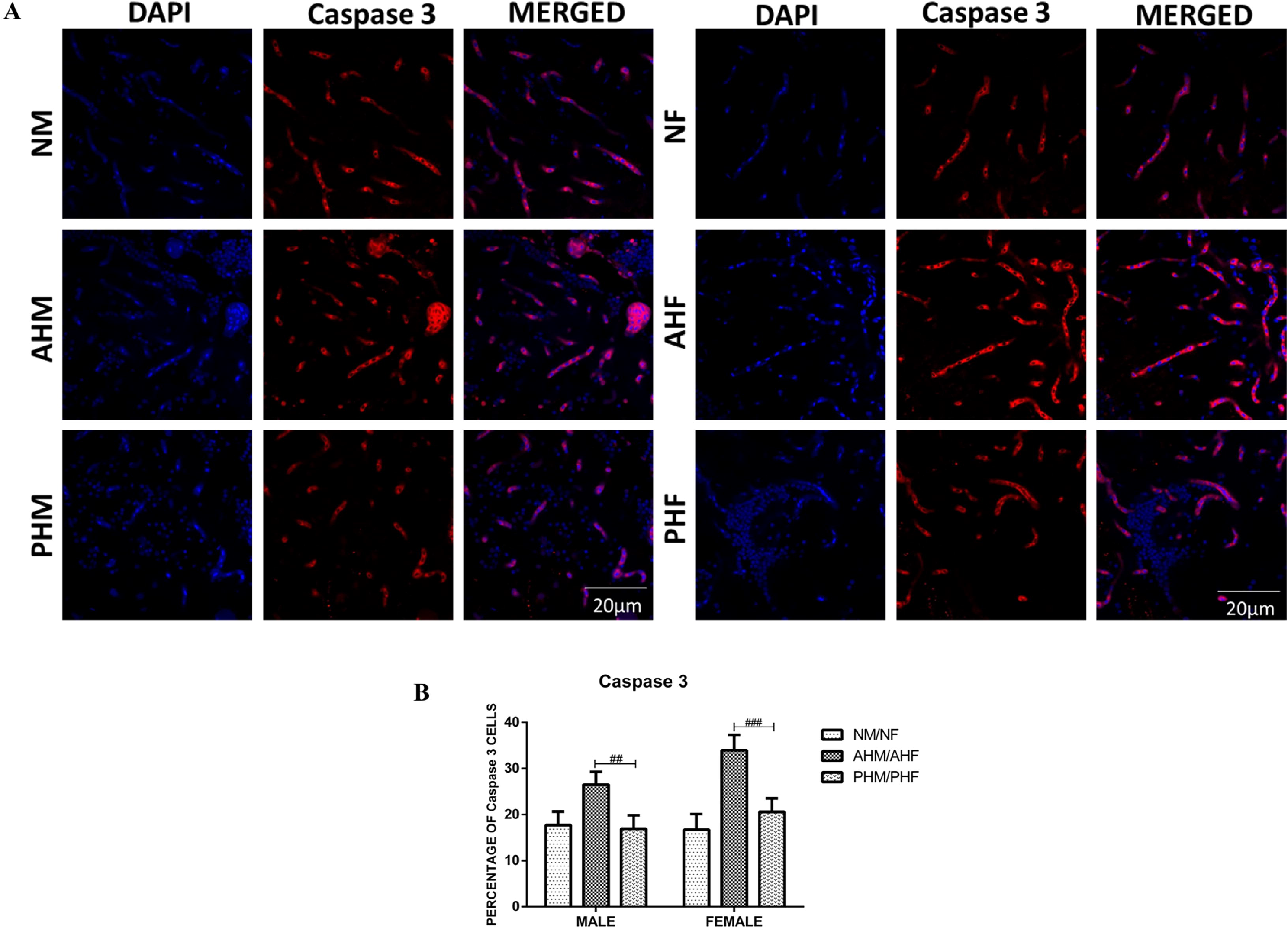
Sex difference in cell death in preconditioning hypoxia vs acute hypoxia. Representative confocal images (60X) of zebrafish male and female brain sections (30μ) showing Cleaved Caspase 3 (red) DAPI (blue) cells (A). Bar indicates the percentage of Cleaved Caspase 3 positive cells (B), n=15. The data are expressed as the mean ± SEM, p < 0.0001, * PHM /PHF vs NM /NF and # PHM/PHF vs AHM/AHF.

### 3.5 Assessment of sex difference in neuroprotection through expression of neuroglial proliferative markers in Preconditioned hypoxia vs Acute hypoxia /Normoxia

Using IFC, the expression of radial glial markers GFAP and BLBP were analysed in both optic tectum and telencephalon of zebrafish brain sections (30 μm, n=15) (Fig 5 A-B). Sex difference in GFAP expression were observed in acute hypoxia group, where in male noticeably less number of GFAP positive cells were observed in acute hypoxia (AHM) as compared to controls (NM). But a contrasting pattern was observed in female acute stress group where its expression was found significantly high (AHF) compared with appropriate controls (NF). In both the sexes, preconditioned stress groups (PHM and PHF) displayed greater percentage of GFAP positive cells as compared to both normoxia and AH groups (Fig 5 A). The statistical analysis showed a significant value [F (2, 59) = 5.638; p<0.0057]. However, no sex difference was observed in the pattern of expression of BLBP in any stress group (neither acute nor preconditioning hypoxia group) but the protection was clearly noticed in preconditioning hypoxia groups (PHM and PHF), irrespective of sex as compared to acute hypoxia groups (AHM and AHF), where the damage appears obvious. The two-way ANOVA showed significant values (F (2, 30) = 18.51; p < 0.0001) (Fig 5 B).

**Fig. 5:**
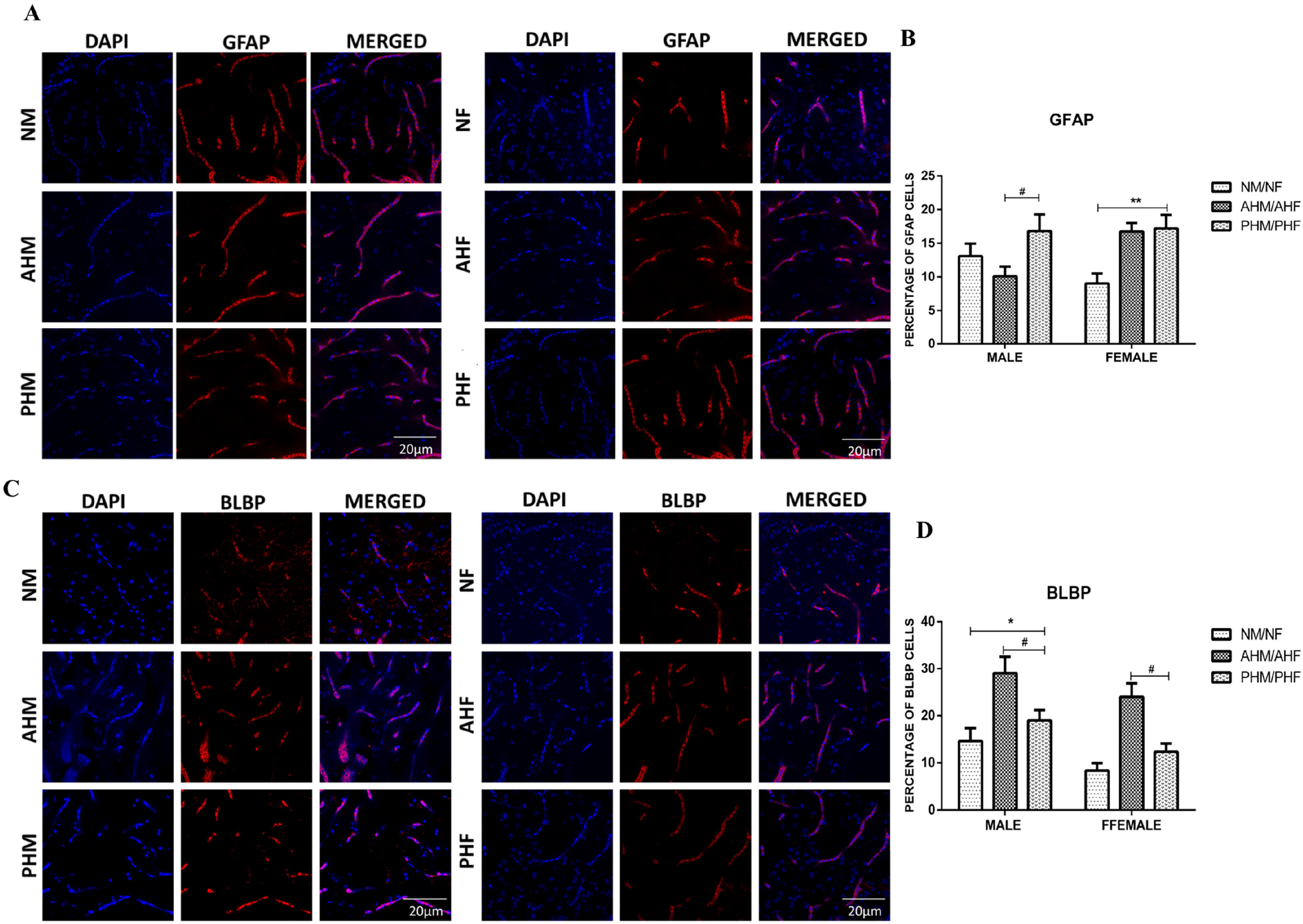
Sex difference in neuroprotection through preconditioning hypoxia. Representative confocal images (60X) of zebrafish male and female brain sections (30μ) showing Gfap and Blbp (red) DAPI (blue) cells (A-B). Bar indicates the percentage of Gfap and Blbp positive cells (CD), n=15. The data are expressed as the mean ± SEM, p < 0.0001, * PHM /PHF vs NM /NF and # PHM/PHF vs AHM/AHF.

Expression of S100B, an astrocytic marker for neural distress and neuropathological conditions, could not show any sex difference in stress regulation pattern, but definitely showed a uniform neuroadaptation or protection in preconditioning hypoxia group (Fig 6). As compared to normoxia, acute hypoxia group showed an upregulation in S100B expression which is not seen in preconditioning hypoxia group. The two-way ANOVA analysis showed a significant value [F (2, 59) = 14.56; p < 0.0001] (n=15).

**Fig. 6:**
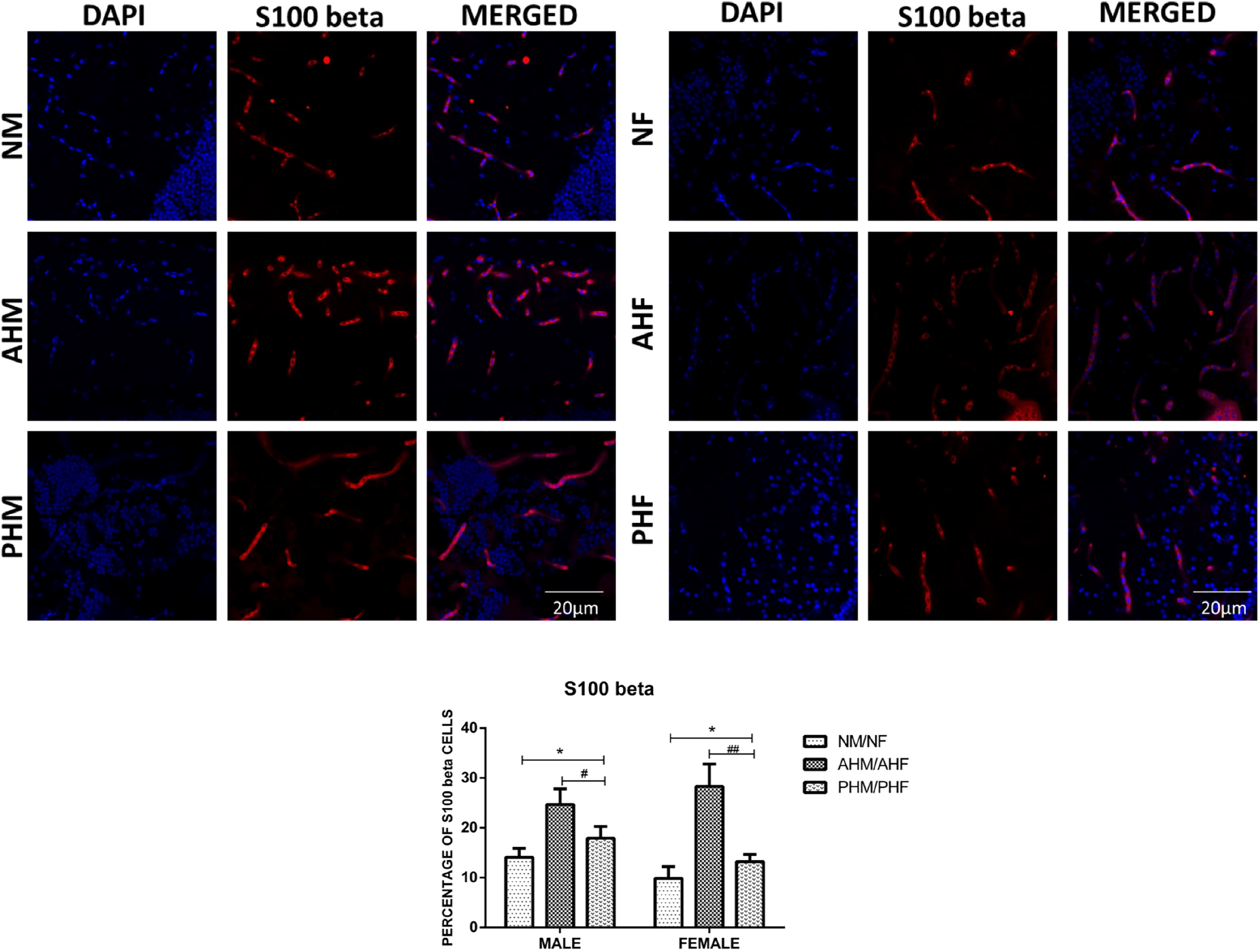
Sex difference in neuroprotection through astrocytic marker. Representative confocal images (60X) of zebrafish male and female brain sections (30 μ) showing S100 B (red) and DAPI (blue) cells (A and C). Bar indicates the percentage of S100B positive cells (B and D), n=15. The data are expressed as the mean ± SEM, p < 0.0001.* PHM /PHF vs NM /NF and # PHM/PHF vs AHM/AHF.

### 3.6 Probing sex difference in preconditioning induced neuroprotection through expression of neural differentiation markers in Acute hypoxia

The expression of NeuN, a marker for mature healthy neuron, showed a sex disparity in acute hypoxia group (Fig 7 A) (n=15). In male acute hypoxia group (AHM) the NeuN positive cells were found considerably less in number than normoxia male (NM), whereas in female (AHF) such change was not found. However, in both preconditioning groups (PHM and PHF) the number of NeuN positive cells was greater than normoxia and acute hypoxia group. The twoway ANOVA analysis showed a significant value [F (2, 59) = 4.572; p < 0.0143].

**Fig. 7:**
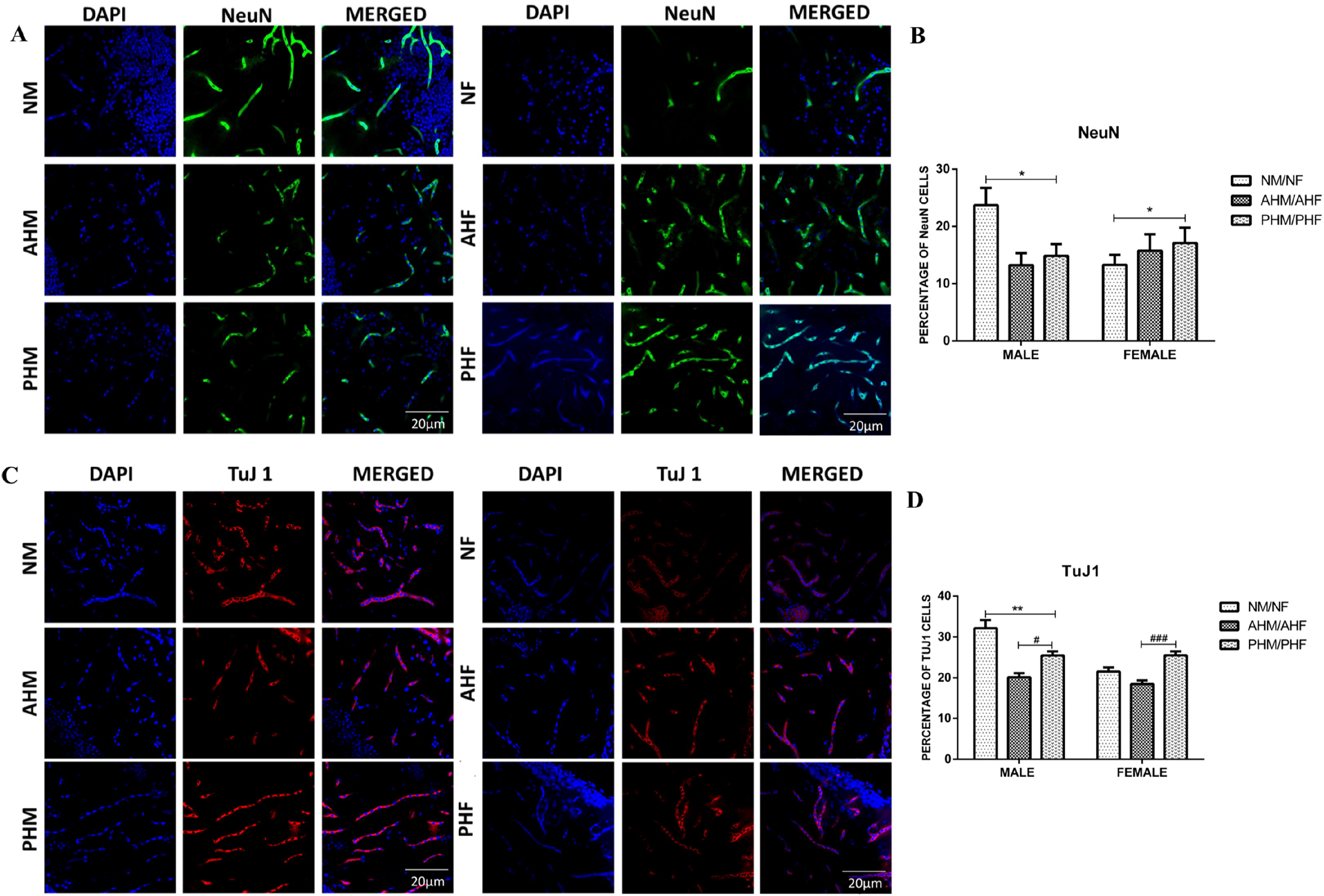
Sex difference in neuroprotection through neural differentiation marker. Representative confocal images (60X) of zebrafish male and female brain sections (30 μ) showing NeuN (green) and Tuj1 (red) DAPI (blue) cells (A and C). Bar indicates the percentage of NeuN and Tuj1 positive cells (B and D), n=15. The data are expressed as the mean ± SEM, p < 0.0001, * PHM /PHF vs NM /NF and # PHM/PHF vs AHM/AHF.

The expression of an early neuron differentiation marker Neuron-specific Class III β-tubulin (TuJ1) showed no sex difference (Fig 7 B) (n=15). Irrespective of the sex, in both the acute stressed groups (AHM and AHF) the number of Tuj1 positive cells were found noticeably less than control groups (NM and NF), whereas the expression of Tuj1 in the preconditioning stressed groups (PHM and PHF) was found to be remarkably more than AHM and AHF. The two-way ANOVA showed a significant value [F (2, 107) = 22.92; p < 0.0001].

### 3.7 Assessing sex difference in neuroprotection through mRNA expression of neurotrophic factors

To find out the role of neurotrophic factors in neuroprotection in preconditioning hypoxia group, we mapped the level of BDNF, NGF and NT3 mRNAs by RT-qPCR. There was a clear upregulation observed in *bdnf* and *ngf* expression in preconditioning hypoxia groups; however, the expression level of nt3 did not show such changes. Interestingly, female showed several folds high expressions in *bdnf* and *ngf* than male preconditioning group (Fig 8).

**Fig. 8:**
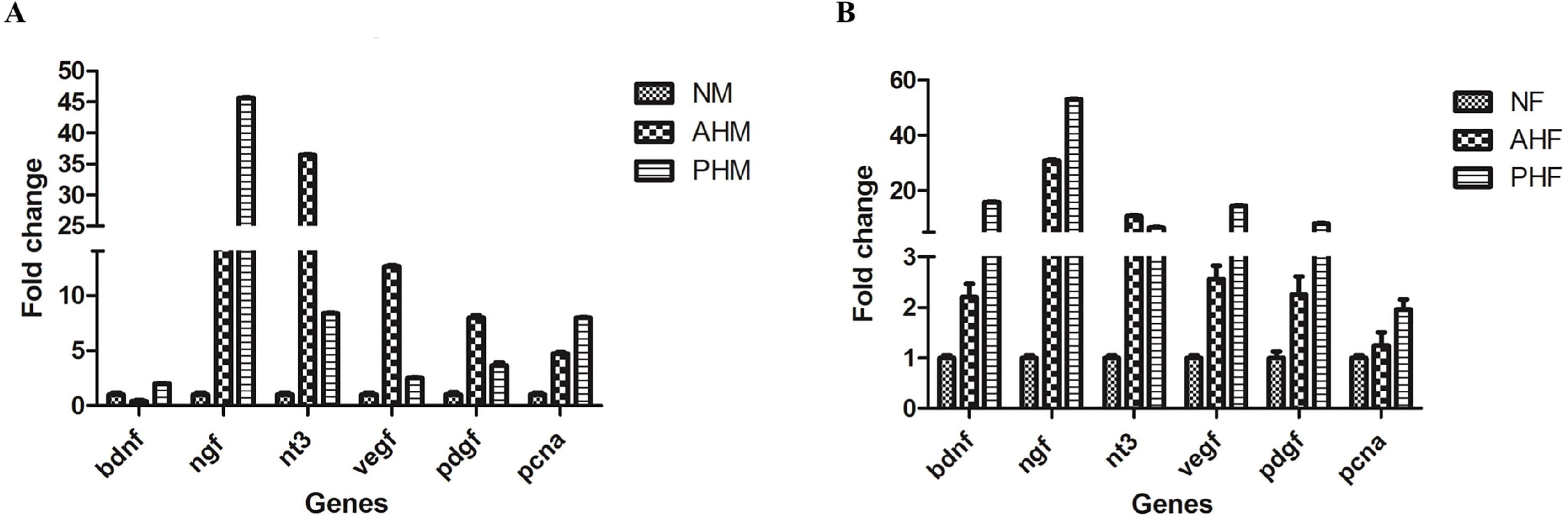
Sex difference neuroprotection through neurotrophic growth factors. mRNA expression of neurotrophic growth factors in NM.AHM, PHM (A) and NF, AHF, PHF (B). (n=4 brains pooled in each group). Here p<0.0001for all the genes when compared with NM/NF vs AHM/AHF and PHM/PHF. Data were analysed by Two-way ANOVA followed by Tukey’s post hoc analysis.

For cell survival study, one of the growth factors (pdgf), angiogenic markers (vegf), and proliferation markers (pcna) were examined at mRNA expression level. Here we could find an increased expression level of vegf and pdgf in AH of both sexes where as in PH male expression of these markers has reduced as compared to AH and in female PH group expression was more as compared to AH. The expression of pcna though did not show any sex difference in expression but an increased expression was seen in PH as compared to AH of both sexes. Though the expression trend was similar (increased) in both sexes still difference in the fold of expression was observed (in male ~5-7 fold where as in female~1.5-2 fold only) [Fig 8, (n=4 brains pooled in each group)].

## 4 DISCUSSION

Improved tolerance of tissues, organs, and even organisms to hypoxic or ischemic stress has been shown to be achieved through the sublethal hypoxic or ischemic exposures (Li *et al*., 2017), the phenomenon of HP is well established in heart (Kloner and Jennings, 2001) and brain (Lu *et al*., 1999b; Lu *et al*., 2005). Repetitive mild hypoxia results in the development of brain HT and cross-tolerance to injurious factors. Such preconditioning acts as a “warning” signal which allows an organism to prepare to cope up with the subsequent more harmful insults. The internal acquired defence mechanisms of hypoxic preconditioning or brain tolerance, are based on genes on mechanisms of adaptation and neuroprotection evolutionary (Michiels, 2004).

Compared to the ischemic preconditioning (by surgical vessel occlusion method), effects and mechanisms of preconditioning by intermittent mild or moderate hypoxia (non-surgically induced) are not well studied; to our knowledge this paradigm of ours first time illustrates a convenient method of hypoxia administration to animals since it does not require surgery. In the current study we have used zebrafish as an alternate model for the preconditioning hypoxia (PH) research. We especially tried to investigate if there is any sex difference in neuroprotection through PH in zebrafish, following the hypothesis that hypoxia preconditioning may give better protection if a sudden severe hypoxic/ischemic insult occurs later. For acute hypoxia (0.6 mg/l) was followed as reported earlier by us (Das *et al*., 2019). Here, equal number of male and female zebrafish were used in each experimental group (normoxia, acute hypoxia and preconditioning hypoxia) and unlike other PH protocol in rodents (Lu *et al*., 2005; Pan *et al*., 2014; Perez-Pinzon *et al*., 1996; Perez-Pinzon *et al*., 1997; Schurr *et al*., 1986; Stagliano *et al*., 1999; Stetler *et al*., 2014; Thompson *et al*., 2013; Yun *et al*., 2014) we have followed a novel protocol for mimicking mini strokes in human. The animals were exposed to mild hypoxic condition intermittently (unpredictable hypoxia stress once or twice in a week for a period of three months) and final day before sacrificing the animals in PH group were subjected to severe acute hypoxia, along with a fresh AH group (without preconditioning) for comparing PH with AH. This method is in contrast to many *in vivo* models of hypoxia that were developed earlier to explore neuroprotection by preconditioning from the point of view of neuro-behaviour, neurophysiology, neurochemistry, neuromorphology, and molecular biology, however in most of the reports preconditioning process was restricted to short duration only prior to acute hypoxia and conducted only in male models. In the vantage point of looking for sex difference in neuroadaptation/ neuroprotection through PH, for the first time we have attempted a novel PH model in zebrafish with intermittent hypoxic episodes for three months before subjecting the animals to a robust hypoxic episode and explored the sex difference in the underlying molecular mechanisms.

Earlier studies using various rodent models have reported that hypoxia causes a number of functional deficits including slowing of various developmental milestones (eye opening, righting reflex), locomotor activity and sensorimotor and memory deficits (Balduini *et al.*, 2000; Lubics *et al*., 2005; McAuliffe *et al*., 2006; Young *et al*., 1986). Though in our earlier report on zebrafish we could show sex difference in the effect of acute hypoxia on neuro behaviour in NTT (Das *et al*., 2019), in the present study we aimed to reveal if there is any sex difference in neuroprotection/neuroadaptation to acute hypoxic insult if the animals had experienced intermittent hypoxic preconditioning. This hypothesis was based on the earlier reports where neuroprotection and improvement in memory after preconditioning (PH) were shown in rodent (Jones *et al*., 2008; Kim *et al*., 2019; Manukhina *et al*., 2020) and zebrafish (Kim *et al*., 2019) models. Similar to those reports, in our study on zebrafish using the novel paradigm of PH remarkable neuroprotection was observed in the preconditioning group upon inducing severe acute hypoxia, when compared with the acute hypoxic group that was not exposed to preconditioning. Moreover, the effect was found to be uniform in both the sexes, without showing any sex difference.

Hypoxia preconditioning protects cells against induced apoptosis, as reported earlier in corneal stromal cells (Xing *et al*., 2006), where a significant reduction in DNA damage was observed in hypoxia preconditioned group. Similarly, in the present study an obvious difference in decrease in DNA damage was observed in PH group as compared to AH group of animals, which again supported the neuroprotective ability of the hypoxic preconditioning. In addition, this study also describes for the first time the sex difference in neuroprotection against the acute hypoxic insult with preconditioning. Male zebrafish brain upon hypoxia insult showed significantly more DNA damage by measure of TUNEL positive cells as compared to the female brain even though both showed neuroprotection by preconditioned hypoxia.

To explore more into the neuroprotective mechanism(s) of hypoxia preconditioning, and into the sex difference, high throughput proteomics on brain samples was performed. The canonical pathways that were revealed by IPA suggested distinct sex difference in one of the top canonical pathways involved. Though EIF2 signalling, mitochondrial dysfunction, sirtuin signalling and synaptogenesis signalling pathways were found common in both male and female brain, these pathways were earlier reported to be affected in various research reports on hypoxia (Ivanova *et al.*, 2018; Koumenis *et al*., 2002; Liu *et al*., 2006; Tafani *et al*., 2016; Zwaans and Lombard, 2014). In order to find out sex difference through our model of PH we have detected the presence of “oxidative phosphorylation pathway” in top canonical pathways of male where as it was not evident in female. Oxidative phosphorylation is the major metabolic pathway for ATP production and the demand for ATP is more in hypoxic condition. As in female we could not observe “oxidative phosphorylation pathway” among the top five canonical pathways, may conferred early recovery from hypoxia in female as compared to male. Regulation of eIF4 and p70S6K signalling pathway was in top five canonical pathways in only female which usually plays a critical role in translational regulation. Translation is the most energy-consuming processes in the cell and a key step in the regulation of gene expression. The eukaryotic translation initiation factor 4E (eIF4E)-binding proteins (4E-BPs) and 70-kDa ribosomal S6 kinases (S6Ks) are few of the mTORC1 substrates which are usually involved in the regulation of mRNA translation. In response to various stimuli, multiple signalling pathways converge on the translational machinery to regulate its function (Roux and Topisirovic, 2018). The early activation of translation machinery upon hypoxia in female only, again gives strength to believe that in female recovery and neural adaptation/ protection is observed faster than male. The most interesting thing was observed in the pattern of expression in canonical pathways compared between both the hypoxic groups with non-stressed normal male (AHM vs NM and PHM vs NM) that it showed a uniformity in pattern of protein expressions. Unlike male, in female three top canonical pathways showed a differentially expressed pattern when compared between acute hypoxia and preconditioning groups with normoxia (AHF vs NF and PHF vs NF). In the present study, since LXR/RXR pathway showed an upregulation in AHF and down regulation in PHF and the activation of LXR/RXR leads to alleviate inflammation (Kong *et al*., 2019), it is legitimate to state that the neuroprotection in female might come faster by reducing inflammation in PHF eventually. Acetyl-CoA, as a central metabolic intermediate, is widely used in macromolecule biosynthesis and energy production to support cell growth and proliferation (Gao *et al*., 2016). In the present study, acetyl-CoA biosynthesis pathway showed an upregulation in PHF and down regulation in AHF which again conferred the better cell growth and proliferation in PHF. The sphingosine kinase-1/sphingosine 1-phosphate (SphK1/S1P) signalling pathway inhibition might indicate a futuristic alternative therapeutic strategy to current mTOR- or VEGF-targeted agents, as S1P is widely appreciated as a general growth-like factor and a potent protector against apoptosis (Bouquerel *et al*., 2016). In the present study an observed upregulation in SphK1/S1P pathway in PHF again supported the notion of neural protection in PH.

Further studying cell death, we have used one of the apoptotic markers cleaved caspase 3 in zebrafish brain section. The result validated the same pattern of neuro-protection through PH. Decrease in cleaved caspase 3 expression in PH in mouse was earlier reported by other research group (Xu *et al*., 2016).

For studying the neuro protective effect of PH we have analysed several neuronal markers for mapping neural proliferation, migration and differentiation. Glial fibrillary acidic protein (GFAP) staining in zebrafish brain section showed an upregulation of expression in PH as compared to normoxia and AH group, which is in contrast to what was reported in other PH model in neonatal rat where there was attenuation in GFAP expression (Chen *et al*., 2015). It is pertinent to mention here that the protocol of preconditioning used in the current study is completely different than other published protocols. In the present study we have observed a sex difference in the expression of GFAP in AH group, as shown by us earlier (Das *et al*., 2019), and speculate to have a role in recovery mechanism after AH where female had shown the recovery faster than male after hypoxia insult. The activation of glial marker in our study can be hypothesized as an adaptive mechanism to tolerate severe hypoxia.

Brain lipid binding protein (Blbp) expression has been shown to be increased in hypoxic conditioning in neural stem cells (Vecera *et al*., 2018). Similarly, here also Blbp expression was found to be upregulated only in acute hypoxic group (AH) when compared with normoxia and preconditioned (PH) groups. It seems that with sudden onset of hypoxia Blbp gets upregulated but as the PH group is already adapted to hypoxic stress, the preconditioning group of animals showed a moderate level of Blbp expression similar to that of normoxia group, which supports neuroprotection through PH.

Similar way we have checked the expression of one of the astrocyte marker S100βwhich has been reported to be increased with acute hypoxia in sheep model (Giussani *et al*., 2005). We too observed an overall increase in its expression in AH group, irrespective of sex. Simultaneously, in PH group moderately low expression of S100β was observed as compared to AH group, though PH group was also treated with similar AH stress, which re-established the neuroprotection phenomenon of preconditioning.

Neuronal nuclear antigen (NeuN) is a nuclear protein known to be widely expressed in mature post mitotic neurons. Hypoxia insult results in loss of NeuN expression as reported in many models (Galle and Jones, 2013; Lavezzi *et al*., 2013). In the present study we could observe a marked loss of NeuN expression in male hypoxic groups (AHM and PHM) but surprisingly in female groups (AHF and PHF) a completely different scenario was seen; a noticeable increase of NeuN expression was observed after severe hypoxia treatment. The sex-specific difference in the expression pattern of NeuN can be explained as either female are less affected by hypoxia insult or they might have better protective mechanism for faster recovery as compared to male.

The expression of neuron-specific class III β-tubulin (TuJ1), a neural differentiation marker, gets enriched in absence of transcription factor HIF1α, as reported in neural stem cell and mouse model (Bohuslavova *et al*., 2019; Vecera *et al*., 2018). Here in our study we have observed a loss of Tuj1 expression in both acute hypoxia groups (AHM and AHF) as compared to the corresponding normoxia groups, which confers with the onset of hypoxia in AH groups Hif 1 α expression is more as reported earlier (Das *et al*., 2019) which resulted in loss of Tuj1 expression, irrespective of sex. But in preconditioning (PH) groups since the system was already adapted to hypoxia and under the protective effect, Hif 1 alpha expression would be less than that in AH resulting in increase of Tuj1 expression as compared to AH.

Repetitive acute intermittent hypoxia (rAIH) increases growth/trophic factor expression in respiratory motor neurons, thereby eliciting spinal respiratory motor plasticity and/or neuroprotection (Satriotomo *et al*., 2016). In our model of PH too, we have found similar pattern of increase in neurotrophic growth factors. BDNF belongs to the neurotrophin family, whose other members are nerve growth factor (NGF) and neurotrophin (NT) 3 (Lu *et al*., 1999a).In the adult zebrafish brain, the localization of *bdnf* transcripts is distributed in all regions of the brain without significant differences between sex,(Cacialli *et al*., 2016) although sex hormones have been recognized to influence the expression and activity of bdnf in mouse stress model (Karisetty *et al*., 2017), where in intact female mouse mRNA expression of bdnf was more as compared to male. Here in our zebrafish hypoxia model also we have observed similar pattern of sex difference in BDNF expression. The expression of pcna which has also role in neurogenesis is upregulated in PH group as compared to AH and normoxia group in accordance with the previous studies on neuronal culture from rat forebrain (Bossenmeyer-Pourie *et al*., 2000). Though expression of pcna is upregulated in both the sexes, still level of expression in male is more as compared to female indicating demand of more neurogenesis in male as neuronal damage is also more.

In most of the experiments in the present study, the neuroprotection was evidently observed in preconditioning groups (PH). It apparently also showed sex difference where female seems to be having a better ability to adapt to PH resulting in faster neural adaptation, in comparison with male.

## 5. CONCLUSION

The present study on preconditioning hypoxia describes the establishment of a protocol in zebrafish to mimic mini strokes or transient ischemic attacks (TIAs) in people. It basically occurs when part of the brain experiences a temporary blockage of blood flow (ischemic attack) resulting in hypoxia. This induces stroke-like conditions that usually resolve within 24 hours. Unlike a major stroke, a mini stroke on its own doesn’t cause permanent disabilities. Hypothesizing that mini stroke events may have similar neuroprotective effect as preconditioning hypoxia does, in the present study for the first time we have followed an intermittent acute hypoxia paradigm for three months, unlike any other published protocol. The aim of the study was to find out whether mini stroke events can protect better from the major stroke event and tried to investigate not only the underlying molecular mechanism/s but also if the neuroprotection process has any sex-specific pattern. Overall, we have observed an obvious neuroprotection through preconditioning hypoxia, irrespective of sex. With few of the neuronal markers (GFAP and NeuN) which showed a distinct sex-specific expression upon hypoxia, it can be concluded that the female seems to be having a better ability to adapt to preconditioning hypoxia to show eventually faster neural adaptation on major hypoxic insult, in comparison with male. A high throughput neural proteome analysis on all the three groups (normoxia, AH, PH), including the sex difference study gives insight not only on the general mechanism underlying the protective effect of mini stroke-like preconditioning, but also shows the possibility of a gender-specific strategy to add in futuristic stroke therapeutics.

## Acknowledgements

This research was supported by the Council of Scientific and Industrial Research (CSIR), India network project (BSC0103-UNDO to SC and AK) and Department of Biotechnology, Government of India (BT/PR14338/MED/30/495/2010 to SC). We thank Director, CSIR-IICT for overall support and KIM department to generate an institutional publication number (IICT/Pubs./2019/437).

